# Identifying Human Specific Adverse Outcome Pathways of Per- and Polyfluoroalkyl Substances Using Liver-Chimeric Humanized Mice

**DOI:** 10.1101/2023.02.01.526711

**Authors:** Dakota R. Robarts, Diego Paine-Cabrera, Manasi Kotulkar, Kaitlyn K. Venneman, Sumedha Gunewardena, J. Christopher Corton, Christopher Lau, Lander Foquet, Greg Bial, Udayan Apte

**Author notes:** **Corresponding Author**: Udayan Apte, PhD, DABT, Department of Pharmacology, Toxicology, and Therapeutics University of Kansas Medical Center, 3901 Rainbow Blvd., MS1018, Kansas City, KS 66160.

## Abstract

**Background:** Per- and polyfluoroalkyl substances (PFAS) are persistent organic pollutants with myriad adverse effects. While perfluorooctanoic acid (PFOA) and perfluorooctane sulfonic acid (PFOS) are the most common contaminants, levels of replacement PFAS, such as perfluoro-2-methyl-3-oxahexanoic acid (GenX), are increasing. In rodents, PFOA, PFOS, and GenX have several adverse effects on the liver, including nonalcoholic fatty liver disease.

**Objective:** We aimed to determine human-relevant mechanisms of PFAS induced adverse hepatic effects using FRG liver-chimeric humanized mice with livers repopulated with functional human hepatocytes.

**Methods:** Male humanized mice were treated with 0.067 mg/L of PFOA, 0.145 mg/L of PFOS, or 1 mg/L of GenX in drinking water for 28 days. Liver and serum were collected for pathology and clinical chemistry, respectively. RNA-sequencing coupled with pathway analysis was used to determine molecular mechanisms.

**Results:** PFOS caused a significant decrease in total serum cholesterol and LDL/VLDL, whereas GenX caused a significant elevation in LDL/VLDL with no change in total cholesterol and HDL. PFOA had no significant changes in serum LDL/VLDL and total cholesterol. All three PFAS induced significant hepatocyte proliferation. RNA-sequencing with alignment to the human genome showed a total of 240, 162, and 619 differentially expressed genes after PFOA, PFOS, and GenX exposure, respectively. Upstream regulator analysis revealed inhibition of NR1D1, a transcriptional repressor important in circadian rhythm, as the major common molecular change in all PFAS treatments. PFAS treated mice had significant nuclear localization of NR1D1. *In silico* modeling showed PFOA, PFOS, and GenX potentially interact with the DNA-binding domain of NR1D1.

**Discussion:** These data implicate PFAS in circadian rhythm disruption via inhibition of NR1D1. These studies show that FRG humanized mice are a useful tool for studying the adverse outcome pathways of environmental pollutants on human hepatocytes in situ.

## Introduction

Per- and polyfluoroalkyl substances (PFAS) are a class of anthropogenic compounds used in a variety of products (Lindstrom et al. 2011). The stability of the fluorinated carbon backbone of these compounds gives them extraordinary water and stain resistance but also makes them non-biodegradable, leading to environmental persistence (Kurwadkar et al. 2022; Rahman et al. 2014). PFAS environmental contamination is of significant concern because of the numerous adverse health effects induced by these chemicals. PFOA and PFOS are the two most abundant PFAS found in the environment (Domingo and Nadal 2019). In humans, both PFOA and PFOS have long half-lives and show substantial bioaccumulation (Fu et al. 2016; Li et al. 2018; Xu et al. 2020). While both PFOA and PFOS have been phased out of production in the United States (US), they are still being imported into the US (Brennan et al. 2021; Sunderland et al. 2019). In recent times, the use of new ‘replacement’ short-chain PFAS with shorter half-lives and lower bioaccumulation such as perfluoro-2-methyl-3-oxahexanoic acid (GenX) has increased (Sunderland et al. 2019). Our recent studies have shown that GenX is potentially hepatotoxic to humans (Robarts et al. 2022b), consistent with previous rodent studies (Chappell et al. 2020; Guo et al. 2021). GenX has been detected in the Cape Fear River Basin in North Carolina and the Rhine River in the Netherlands (Gebbink and van Leeuwen 2020; Guillette et al. 2020; Hopkins et al. 2018; Moller et al. 2010). These rivers are sources of drinking water for multiple cities, increasing the possibility of human exposure.

Human epidemiology studies show that PFAS exposure is associated with adverse hepatic effects, including hypercholesterolemia, changes in bile acid composition, and promotion of non-alcoholic fatty liver disease (NAFLD) (Fenton et al. 2021; Sen et al. 2022; Steenland et al. 2009). The majority of studies identifying PFAS-induced adverse outcome pathways (AOPs) in the liver have been carried out in rodents. These studies suggest that activation of the nuclear receptor peroxisome proliferator-activated receptor alpha (PPARα) by PFOA, PFOS, GenX, and other PFAS is the primary mechanism of action in rodents (Chappell et al. 2020; Rosen et al. 2017). The human relevance of PPARα activation as a mechanism of action underlying hepatocarcinogenicity has been questioned (Corton et al. 2018; Klaunig et al. 2003). Patients who received fenofibrate (a PPARα agonist) treatment for hyperlipidemia do not develop hepatocellular carcinoma (HCC) (Cunningham et al. 2010; Mukherjee et al. 1994). This suggests that PFAS activates different key events (KEs) in rodents compared to humans (Andersen et al. 2021; Corton et al. 2018; Klaunig et al. 2003). There is a critical need to investigate the mechanisms of PFAS-induced hepatotoxicity using more human-relevant models. Previously, other models including transgenic PPARα-knockout mice that express the human gene for PPARα, primary human hepatocytes, and human hepatic spheroids have been used to identify PPARα-independent mechanisms (Beggs et al. 2016; Reardon et al. 2021; Robarts et al. 2022b; Rowan-Carroll et al. 2021; Schlezinger et al. 2021). Here, we utilized a novel *in vivo* human-relevant model to study AOPs in response to PFAS exposure.

The FRG humanized mice are unique in that their livers are repopulated with hepatocytes from a human liver donor (Azuma et al. 2007). Additionally, there is continual genetic selection pressure to maintain the human hepatocytes, preventing mouse cholangiocytes from dedifferentiation into functional mouse hepatocytes. These humanized mice have been utilized as a model to study liver diseases, including NAFLD, viral hepatitis, alcoholic fatty liver disease, and malaria infection (Foquet et al. 2018; Long et al. 2018; Ma et al. 2022; Stone et al. 2021; Tyagi et al. 2018; Wang et al. 2019) as well as a unique human-relevant model to study the effects of chemicals on human hepatocytes *in vivo*. We hypothesize that humanize mice will identify human relevant mechanisms of PFOA, PFOS, and GenX-induced hepatotoxicity.

## Methods

### FRG Humanized Mice

All animal studies were approved and performed under the Institutional Animal Care and Use Committee at the University of Kansas Medical Center (KUMC). FRG KO humanized mice on a NOD background that were 25-week-old were kindly provided by Yecuris, Corp. Humanized mice were generated by injecting cryopreserved human hepatocytes obtained from a single 18-year-old male donor in male triple transgenic male mice (Fah-/-, Rag-/-, and Il2rg-/-) and inducing repopulation of the livers (**Fig. 1A**), as previously described (Azuma et al., 2007). The mice were housed in the KUMC vivarium under a standard 12-hour dark and 12-hour light cycle.

**Figure 1.**
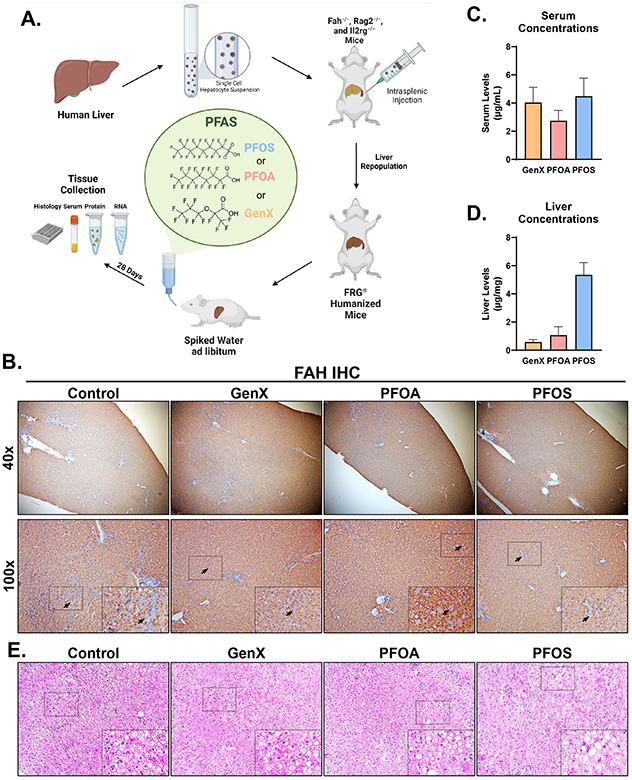
Establishing an *in vivo* human-relevant model to study PFAS. (A) Scheme of the experimental design. (B) Photomicrographs of IHC for FAH in livers of all treatment groups at both 40x and 100x magnifications, where FAH positive hepatocytes indicate human origin. The small box corresponds to the 200x magnification image. Arrowheads are examples of mouse derived hepatocytes. Mass spectrometry analysis of (C) serum and (D) liver PFAS concentrations. The bar represents the mean ± SEM. (E) Liver H&E photomicrographs of all treatment groups at 200x magnification. The small box corresponds to the 400x magnification image.

### PFAS Exposure and Sample Collection

FRG liver-chimeric humanized mice were given water containing either 0.067 mg/L of PFOA, 0.145 mg/L of PFOS, or 1 mg/L GenX ad libitum for 28 days. These concentrations were based on previous reports on occupational exposure of individual PFAS (Beggs et al. 2016; Chang et al. 2014; Olsen et al. 2007). PFOA (Aldrich cat# 77262-50G, lot # BCCB6034), PFOS (Aldrich cat # 77282-10G, lot # BCCC7858), and GenX (Synquest Laboratories cat # 2122-3-09, lot # 00008887) were dissolved in 0.5% Tween-20 at final concentrations of 0.067 g/L, 0.145 g/L, and 1 g/L, respectively. These stocks were diluted to 0.067 mg/L (PFOA), 0.145 mg/L (PFOS), and 1 mg/L (GenX) and supplemented with a final concentration of 0.5% Tween-20 and 3.25% dextrose. Due to that the humanized mice were acclimated to 3.25% dextrose during development, dextrose was added to their drinking water as recommended by Yecuris. The control group was given water containing 0.5% Tween-20 in drinking water supplemented with 3.25% dextrose. All mice were given a tyrosine-free (YF-10^TM^) diet (to prevent liver repopulation with rodent hepatocytes) ad libitum, provided by Yecuris. A sample size of 3 were used in all experimental groups. Mice were euthanized on exposure day 28. Blood was obtained from the retro-orbital sinus, allowed to clot at room temperature for 10 minutes, and then centrifuged at 5000 g for 10 minutes at 4°C to isolate serum. Serum analyses of cholesterol (Fisher cat # 50-489-238), ALT (Fisher cat # 23-666-089), glucose (Fisher cat # 23-666-286), triglycerides (Fisher cat # 23-666-410), free fatty acids (Fisher cat # 50-489-265), and bile acids (Diazyme cat # DZ042A-K01) were performed using kits according to the manufacturer’s protocol. Livers were removed, the gallbladder was separated, and then the liver was weighed to calculate liver weight-to-body weight ratios. A portion of the liver was fixed in 4% formaldehyde for 48 hours, followed by an additional 24 hours in ethanol, and then processed to obtain paraffin-embedded tissue sections for histology. A portion of the liver was cryopreserved in an optimal cutting temperature compound (OCT) for cryosectioning. All remaining liver tissues were stored at −80°C for further analysis.

### PFAS Extraction from Serum and Liver

Serum samples were used with no additional extractions. Liver samples were prepared 3:1 (DI water:tissue mass) and homogenized with a 10/35 PT Polytron homogenizer (Brinkmann Instruments, Westbury, NY). For serum and liver homogenate (25ul) was sub-aliquoted for PFAS extraction, analysis as previously described in depth (Reiner et al. 2009). In brief, proteins were denatured with formic acid, precipitated with cold acetonitrile, and separated by centrifugation (2000 × g for 3 min). Extracted 8 point calibration curves were made using blank rat serum (Pel-Freez Biologicals) spiked with PFAS appropriate for dosed (100 – 2,500 ng/ml) or control (1-100 ng/mL) serum/liver homogenate measurements. Liver homogenate concentrations were corrected for 3:1 dilutions and serum reported as is. An aliquot of the acetonitrile supernatant was placed in an HPLC vial with 2 mM ammonium acetate buffer (pH 6.5) (1:1), and the PFAS concentrations were determined using UPLC–MS/MS.

### Mass Spectrometry Analysis

PFAS analyses were performed using a Thermo Vanquish Horizon ultrahigh performance liquid chromatograph (UPLC) coupled to a Thermo TSQ Quantis triple-quadrupole (QQQ) mass spectrometer operated in negative ion mode. A reversed phase separation of sample components (100 uL) occurred on a Phenomenex Gemini C18, 2 mm × 50 mm, 3.0 μm silica with TMS end-capping column (Torrance, CA) at 55 °C. A Thermo Scientific Hypersil GOLD C18, 1.9 μm, 3 mm × 50 mm was used as a delay column (Waltham, MA) as part of in-house standard practice. The sample was ionized at the mass spectrometer source using electrospray negative ionization. The source and MS/MS parameters were optimized for each analyte individually. Transitions for all ions were observed using multiple reaction monitoring (MRM), and analyte-specific mass spectrometer parameters were optimized for each compound.

### Histology, Immunohistochemistry, and Oil Red O staining

Hematoxylin and eosin staining was performed as previously described on 5 μm thick paraffin imbedded tissue sections (Umbaugh et al. 2022). Paraffin-embedded liver sections (5 μm thick) were used for immunohistochemical analysis of FAH, PCNA, Ki67, NR1D1, CLOCK, and BMAL1 (**Table S1**), as previously described (Robarts et al. 2022a). Flash frozen cryosections (8 μm thick) were used for Oil Red O staining, as previously described (Walesky et al. 2013).

### Protein Isolation and Western Blot Analysis

For each sample, 100 mg of each liver was homogenized in RIPA buffer (Thermo Fisher cat # 89901) containing 1x of phosphatase and protease inhibitors (Thermo Fisher cat # 78427 & 78438). In addition, nuclear, and cytoplasmic fractions were generated using the NE-PER™ Nuclear and Cytoplasmic Extraction kit, according to the manufacturing protocol (Thermo Fisher cat # 78835). Protein concentration was measured using a BCA assay (Thermo Fisher cat # 23225) as previously described (Robarts et al. 2022a). To run the western blots, 100 μg of protein was loaded into each well and run as previously described in depth (Robarts et al. 2022b). The primary antibodies used in this study along with the specific dilution used are shown in **Table S1**. Western blots were imaged and quantified using Image Studio Lite software (Version 5.2). Densitometry was then normalized to the western blot loading control.

### RNA Isolation and qPCR Analysis

RNA was extracted from 50 mg of the liver using Invitrogen™ TRIzol™ Reagent (Thermo Fisher cat # 15596018) as previously described (Robarts et al. 2022a). The RNA was reverse transcribed to cDNA and then utilized for qPCR with 100 ng of cDNA per reaction, as previously described (Apte et al. 2009). The primers utilized are shown in **Table S2**. All genes were normalized to the housekeeping gene (*18S* or *Gapdh*), and fold changes were calculated using the standard 2^−ΔΔCT^ method, as previously described (Livak and Schmittgen 2001).

### RNA-Sequencing

Quality of RNA isolated was assessed using the Agilent TapeStation 4200. All samples had an RNA integrity number equivalent (RIN^e^) value greater than 9.0. cDNA libraries (n of 3 per group) were generated using RNA using the Tecan Universal Plus mRNA-seq kit. These libraries were then sequenced on the NovaSeq 6000 Sequencing System at a sequencing depth of 25 million reads, with 100 cycle base pair paired-end read resolution, provided by the University of Kansas Genomics Core, as previously stated (Gunewardena et al. 2022). An online PFAS dataset using human spheroids as a model was downloaded using SRA-tools from the GEO database, which was analyzed in parallel to the humanized mice RNA-Seq dataset (GSE144775) (Rowan-Carroll et al. 2021). Raw fastq files were then aligned to the human genome (GRCh38), and genes were counted using STAR software (Version 2.3.1u) (Dobin et al. 2013) run on an HPE DL380 Gen10 8SFF CTO high-performance server. The counts were then normalized using the median of ratio method, and differentially expressed gene (DEG) lists were generated using the DESeq2 package (Version 1.28.1) in R Studio (Version 4.0.3, RStudio Team). For the humanized mice study, the samples were compared to the control group. The raw data and normalized data generated were deposited in the GEO database (GSE208636).

### Ingenuity Pathway Analysis

Differentially expressed genes (DEGs) from the humanized mice (p-value < 0.05 and an |foldchange| ≥ 1.5) for each treatment group were uploaded into the Qiagen Ingenuity Pathway Analysis (IPA) software as previously described (Gunewardena et al. 2022; Robarts et al. 2022a). The upstream regulators were then exported from the software using the graphical user interface. Once exported, this dataset was uploaded to RStudio (Version 4.0.3, RStudio Team). The R package ggplot2 (Version 3.3.3) was used to generate dot plots, where size represents the number of genes changed in that pathway, color represents the -log_2_(p-value), and the x-axis illustrates the z-score assigned by the IPA.

### BaseSpace Correlation Engine

Illumina’s BaseSpace Correlation Engine (BSCE, Version 2.0) was used to determine the correlation between the humanized dataset and the human spheroid dataset. Both datasets were uploaded to BSCE using their online interface (https://www.basespace.illumina.com). Briefly, BSCE uses the Running Fisher test to establish positive or negative correlations with corresponding -log(p-values), as previously described in depth (Kupershmidt et al. 2010). These data were imported into RStudio (Version 4.0.3, RStudio Team). The -log(p-values) were assigned a direction; for example, if there was a negative correlation, the -log(p-value) was multiplied by −1. The correlation -log(p-values) for GenX-, PFOA-, PFOS-exposed humanized mice were plotted using the R package ggplot2 (Version 3.3.3) to produce line charts and heatmaps.

### in silico Docking

To determine the interaction of PFOA, PFOS, or GenX with the nuclear receptor NR1D1, we utilized AutoDock Vina (Trott and Olson 2010). The PDB file for NR1D1 (1GA5) was loaded into AutoDock Tools (Version 1.5.6) (Sierk et al. 2001). All ligands and DNA were first deleted from the PDB file. Then, polar hydrogens and Gastereiger charges were added along with the construction of the docking grid (90 × 102 × 110 Å) (Morris et al. 2009). The chemical 3D structures of PFAS were downloaded from PubChem and converted into a PDB format in PyMol (Version 4.6). The PFAS PDBs were prepared using AutoDock Tools (Version 1.5.6) (Adams et al. 2010). The compound was then docked using Vina, as previously described (Akakpo et al. 2019). The interaction of each PFAS with NR1D1 was visualized in PyMol (Version 4.6).

### Graphs and Statistical Analysis

Heatmaps, volcano plots, Venn diagrams, dot plots, and line graphs were produced in R studio (Version 4.0.3, RStudio Team), as previously described using the packages gplots (Version 3.1.1), RcolorBrewer (Version 1,1-2), ggVennDiagram (Version 1.1.1), and ggplot2 (Version 3.3.3) (Gunewardena et al. 2022; Robarts et al. 2022a). Bar graphs and statistical analyses were produced in GraphPad Prism 8. If two groups were being compared, a two-tailed t-test was performed. If three or more groups were compared, an ANOVA was used followed by a Tukey multiple comparison post-hoc test. Statistical significance was considered when the p-value was <0.05.

## Results

### Generating a human-relevant model to study the AOPs of PFAS

To determine the extent of retention of human hepatocytes during the 28-day exposure, we performed immunohistochemistry (IHC) of the enzyme fumarylacetoacetate (FAH), which is expressed only by the transplanted human hepatocytes, on the liver sections of FRG mice (**Fig. 1B**). In all groups, FAH was expressed throughout the entire liver lobule, indicating successful retention and an adequate number of human hepatocytes (**Fig. 1B**). Next, we determined the serum and liver concentrations of PFAS using mass spectrometry. The average serum concentration was 4.04 μg/mL for GenX, 2.74 μg/mL for PFOA, and 4.48 μg/mL for PFOS (**Fig. 1C**). PFAS were not detected in the control mice. This indicates that both PFOA and PFOS levels were in the same magnitude of observed in occupational workers (0.07-5.10 μg/mL and 0.14-3.50 μg/mL, respectively) (Beggs et al. 2016; Chang et al. 2014; Olsen et al. 2007). GenX levels were comparable to those of PFOS. Further, all PFAS accumulated in the liver, with PFOS at the highest concentration of 5.35 μg/mg followed by PFOA at 1.07 μg/mg (**Fig. 1D**). GenX had the lowest amount of hepatic accumulation at 0.58 μg/mg (**Fig. 1D**). Hematoxylin and Eosin (H&E) staining showed mild to moderate steatosis in the livers of all PFAS treated mice with PFOA treatment showing the highest accumulation of fat (**Fig. 1E**).

### PFAS altered LDL/VLDL, bile acids, and lipid deposition in the liver

The liver-weight-to-body-weight ratio did not indicate any PFAS-related changes over the control group (**Fig. 2A**). Serum ALT, a marker of liver injury, and other metabolic markers, including serum glucose, triglycerides, and free fatty acids, showed no significant difference between any treatment groups (**Fig. 2B-E**). Previous studies have shown that serum cholesterol levels are elevated in humans exposed to PFAS (Andersen et al. 2021; Frisbee et al. 2010; Steenland et al. 2009). We measured total cholesterol, LDL/VLDL and HDL in the serum of the humanized mice following PFAS treatment (**Fig. 2F**). Total cholesterol was similar in PFOA and GenX treated mice but was significantly lower in PFOS treated mice. PFOS also caused a significant decrease in LDL/VLDL compared to the control, GenX, and PFOA groups. Furthermore, GenX showed a significant increase in LDL/VLDL compared to the control group (**Fig. 2G**). However, PFAS-induced changes in HDL did not reach statistical significance, although a trend of decreased HDL was noted in the PFOA and GenX treatment group (**Fig. 2H**).

**Figure 2.**
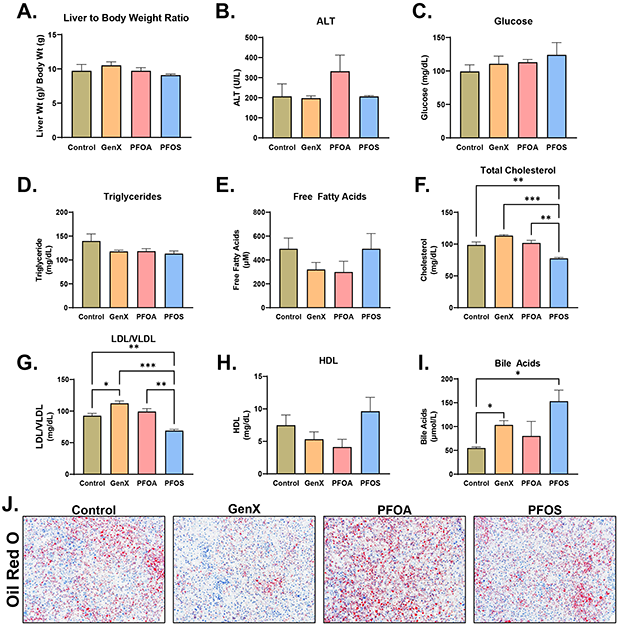
Serum profiles and hepatic lipid deposition of humanized mice exposed to PFAS. (A) Liver weight-to-body weight percentage in all treatment groups. Measurements of ALT, (C) Glucose, (D) Triglycerides, (E) Free Fatty Acids, (F) Total Cholesterol, (G) LDL and VLDL, (H) HDL, and (I) bile acids in the serum of all treatment groups. Bar graphs represent the mean ± SEM, where * p < 0.05, ** p < 0.01 and *** p < 0.001. (J) Photomicrographs of Oil Red O staining in the livers of all treatment groups to indicate lipid deposition.

PFAS exposure in humans has been strongly correlated with alterations in bile acids in the serum (Sen et al. 2022). We measured bile acids in serum of humanized mice exposed to PFOA, PFOS, and GenX and found significantly elevated bile acids in GenX and PFOS treated mice (**Fig. 2I**). We performed Oil Red O staining to visualize lipid accumulation, which showed significant lipid accumulation following PFOA treatment (**Fig. 2J**). Interestingly, GenX had less lipid accumulation compared to the control group, and PFOS showed no difference.

### PFAS induced hepatocyte proliferation in humanized mice after 28 days of exposure

To determine the extent of PPARα in mouse and human derived hepatocyte, species specific qPCR was performed. This showed an induction in mouse PPARα target genes whereas human had no change in gene expression in PFOA, PFOS, and GenX treatments (**Fig. S1A-C)**. To investigate whether PFOA, PFOS, or GenX treatment resulted in the induction of cell proliferation in humanized mice, we performed IHC for PCNA and Ki67. PFOA and PFOS caused a significant induction of cell proliferation compared to the control (**Fig. 3A–C)**, but GenX treatment did not cause significant proliferation. Consistently, qPCR analysis showed significant induction in cyclin D1, the major cell cycle regulator, in PFOA and PFOS treated mice with no change in the GenX treated mice compared to the control (**Fig. 3D**). Further, western blot analysis showed an induction of cyclin D1 and p-Rb proteins in the livers of PFOA, PFOS, and GenX treated mice (**Fig. 3E**).

**Figure 3.**
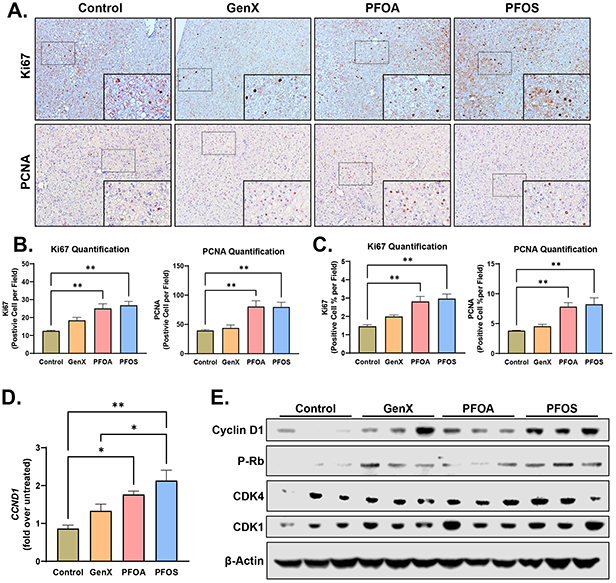
PFAS induced significant proliferation in Humanized Mice after 28 day exposure. (A) Photomicrographs (200X) of Ki67 and PCNA. The small box corresponds to the 400x magnification image. IHC in the livers of all treatment groups with respective quantification at 200x magnifications of positive (B) cell number and (C) cell percentage. (D) Gene expression analysis using qPCR on the proliferative genes Cyclin D1 (*CCND1*). Data were normalized to 18s and then to the control group. Bar graphs represent the mean ± SEM. (E) Western blot analysis of whole liver lysates for the proliferative mitogens cyclin D1, phosphorylated RB, CDK4, and CDK1 with β-actin as a loading control.

### Significant transcriptome changes were exhibited in the livers of humanized mice exposed to PFAS

To determine the global effect of human-relevant PFAS exposures on humanized mice, we performed bulk RNA-sequencing (RNA-Seq). Livers of FRG humanized mice have over 80% human hepatocytes, but all the non-parenchymal cells (NPCs) are of mouse origin. To determine specific changes in human hepatocytes and exclude changes occurring in mouse NPCs, all alignments were performed on the human genome. We compared ligand-binding nuclear receptors from human RNA-Seq datasets to humanized mice RNA-Seq using a rank-based method and found that the top expressed receptors were RXRα, PPARα, and AR, whereas the least expressed receptor was RXRγ (**Fig. S2A**). Cluster analysis shows that transcriptomic changes following PFOA and PFOS were similar to each other but distinct from those induced by GenX (**Fig. S2B**). A global analysis of all differentially expressed genes (DEGs) found that PFOA caused significant downregulation of 154 genes and significant upregulation of 86 genes, with the top altered genes including *KRT23*, *NRN1*, *RGS2*, *MRC2*, and *UCP2* (**Fig. 4A, Table 1**). PFOS treatment downregulated 120 genes and upregulated 42 genes. *HSPA6*, *GRIP2*, *KRT25*, *ECEL1*, and *RASGRP2* were the top altered genes after PFOS treatment (**Fig. 4B, Table 1**). GenX caused the greatest number of gene changes; 446 genes were downregulated, and 173 genes were upregulated (**Fig. 4C**). The top 5 DEGs after GenX treatment were *KIF18A*, *MRC2*, *PLXNA4*, *SLITRK3*, and *ADAMTS1* (**Table 1**). Venn diagrams of DEGs across chemicals showed that GenX uniquely upregulated 119 and uniquely downregulated 310 genes, PFOA caused upregulation of 45 and downregulation of 49 unique DEGs whereas PFOS caused upregulation of 9 and downregulation of 42 unique DEGs (**Fig. 4D–E**). A total of 12 DEGs were commonly upregulated across all PFAS groups, including *COL6A2*, *MYO16*, *LAMC3*, and a variety of metallothionein genes (**Fig. 4D–E, Table 2**). In addition, 23 DEGs were commonly downregulated across all exposures, including *CYP3A7*, *SCL7A10*, *MRC2*, and cyclins A2 and B1 (**Fig. 4D–E, Table 3**). To visualize alterations in cytochrome P450s and phase 2 drug metabolism enzymes (DMEs), heatmaps were made using the fold change values when compared to the control group (**Fig. S2C-B**). *CYP2A7, CYP3A4, CYP2A13, CYP26A1*, and *UGT1A3* were the most commonly induced DMEs (**Fig. S2C–D**).

**Table 1.**
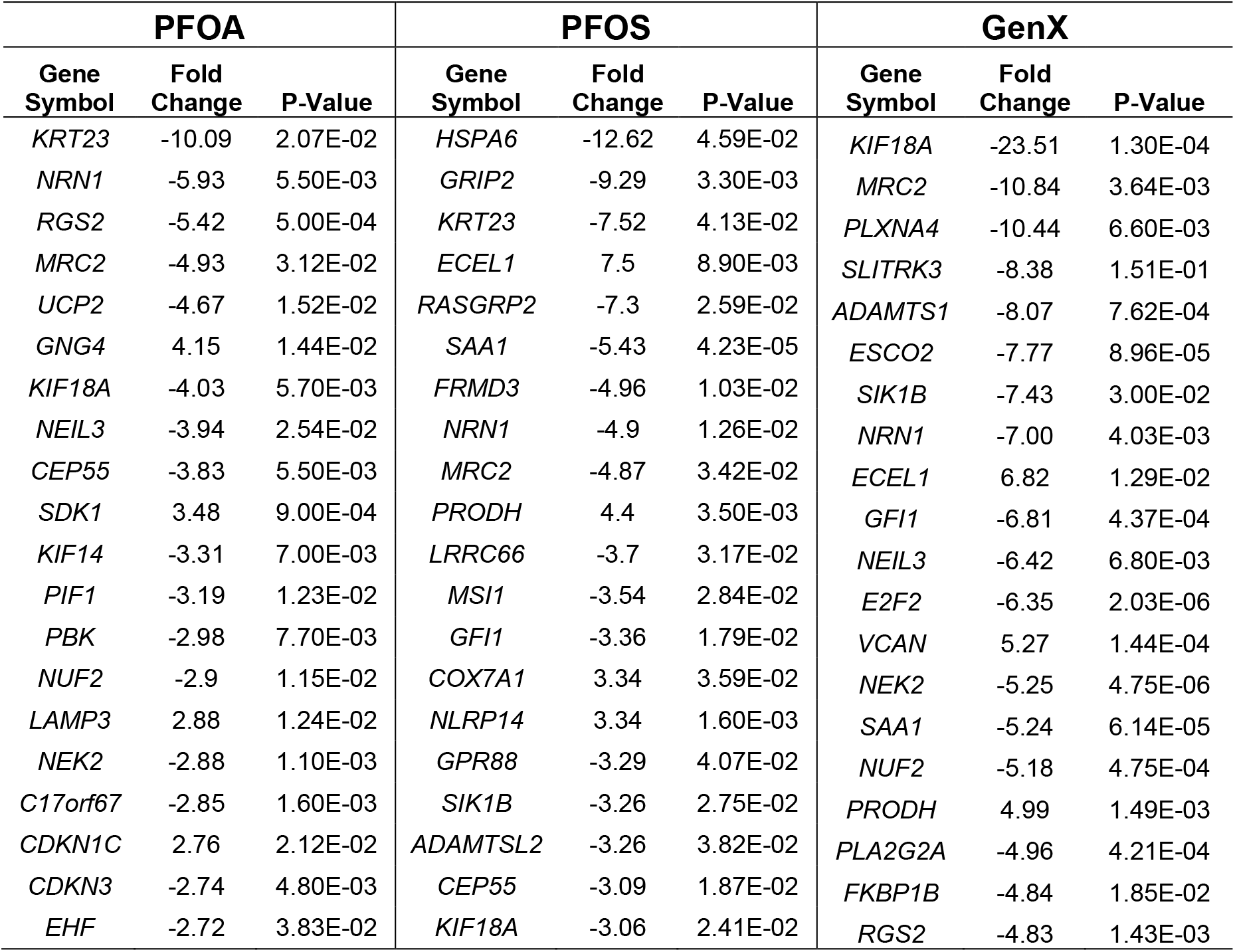
Top 20 |fold change| of DEGs that are significantly altered in livers from PFOA, PFOS, and GenX exposed humanized mice.

**Table 2.** Common upregulated DEGs between PFOS, PFOA, and GenX exposed mice.

|  | **PFOA** |  | **PFOS** |  | **GenX** |  |
| --- | --- | --- | --- | --- | --- | --- |
| **Gene Symbol** | **Fold Change** | **P-Value** | **Fold Change** | **P-Value** | **Fold Change** | **P-Value** |
| *C19orf71* | 1.53 | 2.14E-02 | 1.51 | 2.57E-02 | 1.67 | 5.31E-03 |
| *COL6A2* | 1.68 | 1.85E-02 | 2.04 | 1.24E-03 | 4.02 | 2.62E-10 |
| *LAMC3* | 2.62 | 6.38E-03 | 2.08 | 4.36E-02 | 2.31 | 2.03E-02 |
| *MT1E* | 2.01 | 1.34E-02 | 1.88 | 2.60E-02 | 2.20 | 5.34E-03 |
| *MT1F* | 2.57 | 2.78E-02 | 2.62 | 2.49E-02 | 3.19 | 6.97E-03 |
| *MT1G* | 2.47 | 4.97E-03 | 2.26 | 1.12E-02 | 2.67 | 2.28E-03 |
| *MT1H* | 2.25 | 1.42E-02 | 1.96 | 4.15E-02 | 2.17 | 1.94E-02 |
| *MT1M* | 2.45 | 3.05E-02 | 2.52 | 2.55E-02 | 3.16 | 5.53E-03 |
| *MT1X* | 2.52 | 3.90E-03 | 2.31 | 8.99E-03 | 2.85 | 1.05E-03 |
| *MT2A* | 1.93 | 2.31E-02 | 1.95 | 2.13E-02 | 2.24 | 5.43E-03 |
| *MYO16* | 1.51 | 2.20E-04 | 1.53 | 1.94E-04 | 1.65 | 9.36E-06 |
| *SPTBN4* | 2.30 | 6.37E-03 | 2.55 | 2.27E-03 | 2.47 | 3.25E-03 |

**Table 3.** Common downregulated DEGs between PFOS, PFOA, and GenX exposed mice.

|  | **PFOA** |  | **PFOS** |  | **GenX** |  |
| --- | --- | --- | --- | --- | --- | --- |
| **Gene Symbol** | **Fold Change** | **P-Value** | **Fold Change** | **P-Value** | **Fold Change** | **P-Value** |
| *ADAM19* | -1.77 | 2.38E-02 | -1.84 | 1.70E-02 | -2.58 | 3.02E-04 |
| *ANLN* | -2.20 | 2.11E-03 | -1.73 | 3.18E-02 | -3.66 | 1.59E-06 |
| *BCL2L14* | -2.44 | 3.66E-02 | -2.78 | 2.10E-02 | -2.40 | 4.46E-02 |
| *BIRC5* | -1.76 | 2.00E-02 | -1.92 | 7.80E-03 | -3.17 | 6.14E-06 |
| *CCNA2* | -1.86 | 6.64E-03 | -1.66 | 2.69E-02 | -2.28 | 4.33E-04 |
| *CCNB1* | -1.92 | 5.00E-03 | -1.62 | 3.83E-02 | -3.17 | 2.59E-06 |
| *CDC20* | -2.65 | 1.90E-03 | -1.92 | 3.72E-02 | -4.06 | 1.51E-05 |
| *CENPE* | -2.63 | 1.65E-03 | -2.14 | 1.25E-02 | -2.19 | 1.07E-02 |
| *CYP27B1* | -2.00 | 1.51E-02 | -1.76 | 4.71E-02 | -2.51 | 2.37E-03 |
| *CYP3A7* | -1.82 | 1.94E-02 | -1.84 | 1.75E-02 | -2.26 | 1.59E-03 |
| *DLGAP5* | -1.93 | 2.60E-02 | -1.87 | 3.59E-02 | -3.07 | 4.99E-04 |
| *HSPA5* | -1.54 | 1.57E-04 | -1.60 | 3.70E-05 | -1.88 | 3.20E-08 |
| *JCAD* | -2.23 | 3.64E-02 | -2.21 | 4.19E-02 | -2.18 | 4.57E-02 |
| *KIF18A* | -4.03 | 5.70E-03 | -3.06 | 2.41E-02 | -23.51 | 1.30E-04 |
| *KIF20A* | -1.92 | 3.20E-03 | -1.69 | 1.81E-02 | -3.11 | 1.45E-06 |
| *KPNA2* | -1.86 | 1.22E-05 | -1.55 | 2.23E-03 | -2.34 | 4.77E-09 |
| *MRC2* | -4.93 | 3.12E-02 | -4.87 | 3.42E-02 | -10.84 | 3.64E-03 |
| *NDC80* | -1.86 | 8.72E-03 | -1.99 | 4.82E-03 | -2.49 | 3.14E-04 |
| *NEK2* | -2.88 | 1.06E-03 | -2.01 | 2.69E-02 | -5.25 | 4.75E-06 |
| *NRN1* | -5.93 | 5.46E-03 | -4.90 | 1.26E-02 | -7.00 | 4.03E-03 |
| *RDH12* | -2.28 | 1.74E-02 | -2.15 | 2.81E-02 | -2.21 | 2.32E-02 |
| *SLC7A10* | -2.10 | 6.75E-03 | -2.74 | 3.66E-04 | -2.99 | 1.25E-04 |
| *TIGAR* | -2.29 | 2.56E-03 | -1.89 | 2.06E-02 | -2.24 | 4.01E-03 |

**Figure 4.**
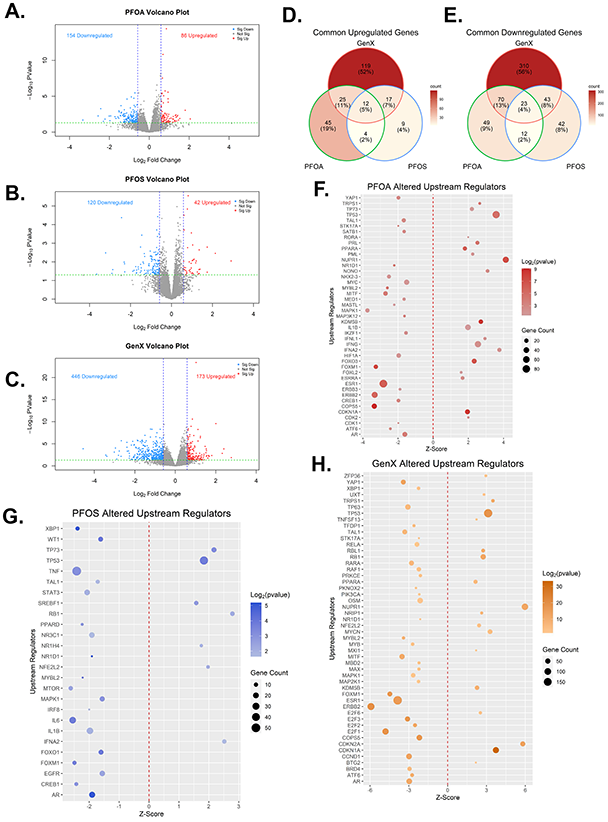
PFAS induces significant transcriptome alterations and upstream regulators. Volcano plots of the DEGs from the (A) PFOA, (B) PFOS, and (C) GenX treated humanized mice, where red and blue dots represent significantly upregulated and downregulated genes, respectively. The blue dashed lines represent a Fold Change of 1.5, and the red dashed line represents a p-value of 0.05. Venn diagrams of DEGs of (D) upregulated and (E) downregulated genes, with shades of red representing the number of DEGs in each segment of the diagram. Dot plot of the IPA upstream analysis for (F) PFOA, (G) PFOS, and (H) GenX. The color represents the log_2_(p-value), the size of the dot represents the number of DEGs in the pathway, the z-score represents the activation status (negative = inhibition and positive = activation), and the red dashed line represents a z-score of 0.

Next, to compare these *in vivo* data to *in vitro* data, we compared the humanized mice DEGs of PFOA and PFOS with an RNA-Seq dataset derived from human hepatic spheroids treated with either PFOA or PFOS at multiple concentrations (PFOA: 0.02, 0.1, 0.2, 1, 2, 10, 20, 50, or 100 μM, and PFOS: 0.02, 0.1, 0.2, 1, 2, 10, or 20 μM) and time points (1-, 4-, 10-, or 14-day) (Rowan-Carroll et al. 2021). We found that our PFOS treated humanized mice were most similar to the 1 μM treatment for 14 days and the most different to the 2 μM treatment for 1 day (**Fig. S3A-B**). The PFOA treated mice were most similar to the 14-day spheroid exposure at the 10 μM concentration (**Fig. S3A, S2C**). Interestingly, we found that GenX was most similar to the PFOA spheroids treated at the 10 μM for 10 days and most dissimilar to the PFOS 1-day exposure at 2 μM (**Fig. S3A**).

To provide insight into mechanisms that were contributing to these DEGs altered in PFOA, PFOS, and GenX exposed mice, we uploaded the DEGs into the Ingenuity Pathway Analysis (IPA) software to determine altered upstream regulators. We found that PFOA significantly inhibited androgen receptor (AR), FOXM1, and NR1D1, and induced activation of TP53, PPARα, and the proinflammatory regulators INFγ and IL1B (**Fig. 4F**). PFOS had the fewest total altered upstream regulators; PFOS significantly inhibited FOXM1, IL6, IL1B, and NR1D1, and activated TP53 and SREBF1 (**Fig. 4G**). GenX had the most significantly altered upstream regulators. GenX inhibited ESR1, RAF1, NR1D1, and TAL1 and activated TP53, PPARα, and RB1 (**Fig. 4H**).

### NR1D1 is significantly disrupted in PFAS exposed humanized mice

An interesting *in vivo* pathway was identified in the IPA analysis. The nuclear receptor NR1D1 (also known as Rev-Erbα) was predicted to be significantly inhibited in PFOA, PFOS, and GenX treated humanized mice (**Fig. 4F-H**). NR1D1 is a nuclear receptor critical for regulating circadian rhythm at the molecular level. Two circadian transcriptional activators, CLOCK, and BMAL1, heterodimerize to activate a plethora of genes, including NR1D1. The upregulation of NR1D1 initiates the negative feedback loop that occurs during the light cycle. Once translated into protein, NR1D1 translocates to the nucleus, binds to the promoter of BMAL1, and represses BMAL1 expression. This causes the intracellular levels of NR1D1 to drop to restart the cycle of regulating circadian rhythm (Tahara and Shibata 2016).

To investigate the disruption of NR1D1, we first performed qPCR on genes regulated by CLOCK and BMAL1, including *NR1D1*, *CRY2*, *PER2*, and *PER1*. *NR1D1*, *CRY2*, and *PER2* were all significantly upregulated in GenX, PFOA, and PFOS exposed humanized mice (**Fig. 5A**). *PER1* was significantly elevated in the PFOA and PFOS exposed mice but not in GenX treated mice (**Fig. 5A**). Western blot analysis showed a significant induction of NR1D1 in all treatment groups compared to the control group, with the strongest induction following GenX treatment (**Fig. 5B–C**). CLOCK protein expression did not change, as expected, because NR1D1 does not regulate CLOCK expression (**Fig. 5B–C**). BMAL1 protein showed no significant changes in all treatment groups (**Fig. 5B–C**), which was consistent with its mRNA expression (**Fig. S4B**). IHC of CLOCK and BMAL1 corroborated the western blot data (**Fig. S4A**). However, IHC of NR1D1 showed significant localization in the nucleus in PFOA, PFOS, and GenX exposed humanized mice with little localization in the control group (**Fig. 5D**). Further, we performed Western blot analysis of NR1D1, CLOCK, and BMAL1 using cytoplasmic and nuclear fractions, which yielded similar results showing no change in CLOCK and BMAL1 and a significant amount of NR1D1 protein in the nucleus (**Fig. 5E**).

**Figure 5.**
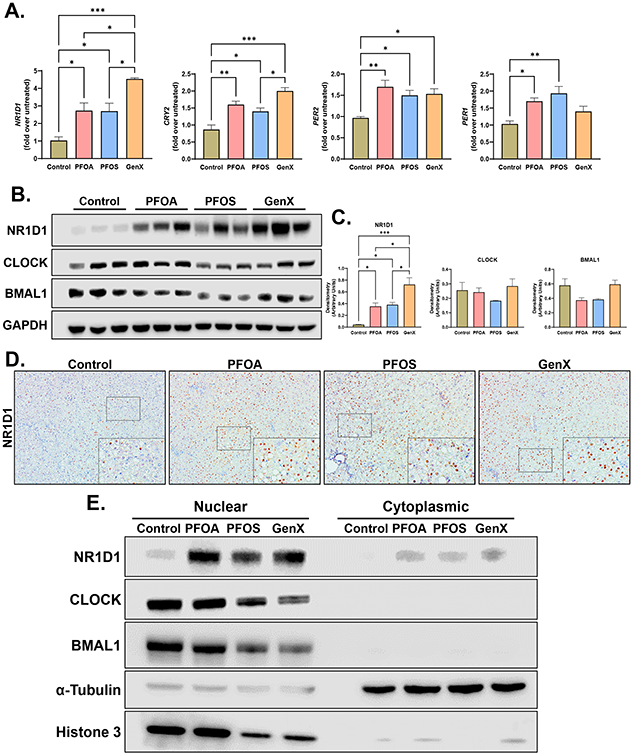
NR1D1 was significantly induced in all PFAS treatment groups. (A) qPCR analysis of livers from PFAS treatment groups of E-box regulated genes, including *NR1D1*, *CRY2*, *PER2*, and *PER1*. Samples were first normalized to the housekeeping gene 18s and then to the control group. (B) Western blot analysis of NR1D1 and the regulators BMAL1 and CLOCK. GAPDH was used as a loading control. (C) Densitometric analysis of the western blots, where the densitometry of each band was normalized to GAPDH. (D) Photomicrographs of liver IHC of NR1D1 from PFAS treated mice at 200x resolution. The small box corresponds to the 400x magnification image. (E) Western blot analysis of nuclear and cytoplasmic fractionation from control and PFAS treated livers of NR1D1, CLOCK, and BMAL1. Histone 3 and α-tubulin were used as loading controls for the nuclear and cytoplasmic fractions, respectively. (F) Densitometry of the fractionated western blots. All bar graphs represent the mean ± SEM. * p < 0.05, ** p < 0.01 and *** p < 0.001

Due to the strong translocation of NR1D1 into the nucleus and the absence of suppressed BMAL1 expression, we examined whether PFAS directly inhibits NR1D1. To do this, we performed *in silico* ligand docking with each PFAS into the crystalized structure of the DNA-binding domain of NR1D1. We found that PFOA, PFOS, and GenX all docked successfully into the DNA-binding pocket, with delta G values of −7.0, −6.5, and −5.4, respectively (**Fig. 6A–C**). Altogether, our data utilizing humanized mice indicate that PFAS inhibit NR1D1-mediated regulation of circadian rhythm genes (**Fig. 6D**).

**Figure 6.**
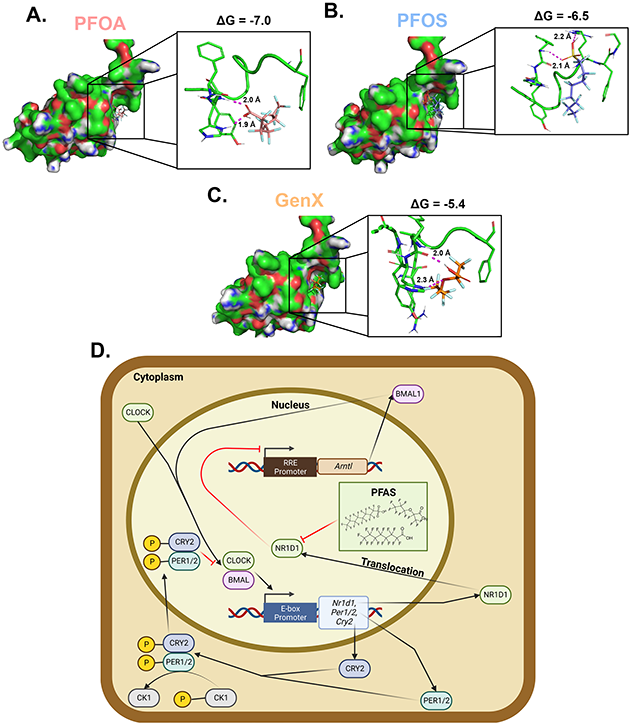
PFAS act as an NR1D1 antagonist by binding to the DNA binding domain. Docking of (B) PFOA, (C) PFOS, and (D) GenX into the crystal structure of NR1D1, with corresponding delta G values and predicted protein-compound interactions. (E) Mechanism of action of PFAS through NR1D1, causing dysregulation of the circadian rhythm E-box pathway.

## Discussion

### A novel model to identify AOPs involved in PFAS toxicity

PFAS such as PFOA, PFOS, and GenX are activators of PPARα the primary mechanism by which PFAS induce hepatotoxicity (Conley et al. 2022; He et al. 2022; Pan et al. 2021; Robarts et al. 2022b; Rosen et al. 2010; Wang et al. 2017; Wolf et al. 2008). In rodents, PPARα activation is the first key event (KE) in the development of hepatotoxicity (Corton et al. 2018). However, in humans, PPARα activation is not thought to be a KE due to lower expression levels of full-length PPARα compared to rodents (Corton et al. 2018; Palmer et al. 1998) and to a lesser extent differences in affinities between species (Keller et al. 1997; Takacs and Abbott 2007). Because of this, studies in *Ppara*-null rodents and *in vitro* models including primary human hepatocytes, organoids, and cell spheroids have been conducted and to identify important human relevant mechanistic information (Beggs et al. 2016; Reardon et al. 2021; Robarts et al. 2022b; Rowan-Carroll et al. 2021; Schlezinger et al. 2021). One issue with *in vitro* models is the lack of cell-cell communication and cross-organ communication that are critical contributors to toxicity mechanisms. In that regard, the humanized-mouse model used in these studies can overcome many of these challenges, because the mice possess human hepatocytes and an intact *in vivo* system, which can potentially reveal more human-relevant AOPs. This was exhibited in which no changes in circadian rhythm were detected in the human spheroid dataset. In addition, PPARα activation in humanized mouse models was less prominent in human hepatocytes compared to rodent hepatocytes utilizing the agonist fenofibrate (de la Rosa Rodriguez et al. 2018). Our study corroborates this with an induction of rodent PPARα target gene expression, while no change in the expression of the orthologous human gene.

### Cholesterol and bile acid alterations induced by PFAS exposure

One of the main findings of epidemiological studies was that PFAS (particularly PFOA and PFOS) exposure has a strong correlation with increasing cholesterol levels in serum (Andersen et al. 2021; Blake and Fenton 2020; Frisbee et al. 2010; Rogers et al. 2021; Rosato et al. 2022; Steenland et al. 2009). Steenland et al. (2009) found that with increasing PFOA and PFOS concentrations, the odds ratio of having higher levels of cholesterol increased, particularly in LDL/VLDL but not HDL. Another study showed that in children PFOA and PFOS were significantly associated with increases in serum total cholesterol and LDL levels (Frisbee et al. 2010). These data suggest that, in humans, PFAS exposure leads to hypercholesterolemia. The effect of GenX exposure on cholesterol levels is unknown. Our data showing a decrease in total cholesterol after PFOS exposure in humanized mice contradict the epidemiology studies. One possibility behind this discrepancy could be that our model exposure window (28 days) was not adequately long enough to induce hypercholesterolemia. When measuring the specific forms of cholesterol, we found that GenX did significantly induce LDL/VLDL with a slight decrease in HDL, indicating that it could potentially affect cholesterol levels in humans. Interestingly, these cholesterol changes in GenX were accompanied by a decrease of lipid accumulation in the liver. This could be attributed to the increased flux of lipids in the form of LDL/VLDL into the serum. Whereas PFOA had an increase in lipid deposition in the liver with no changes in LDL/VLDL but had a trend in decreased HDL. The flux of lipids out of the liver could explain these steatosis phenotypes.

We observed a significant increase in serum bile acids following PFOS and GenX treatments. Bile acids are produced from cholesterol by a series of steps, the first of which is catalyzed by the hepatic CYP7A1 enzyme. However, no significant differences in the expression of *CYP7A1* were observed in PFOA, PFOS, and GenX exposed humanized mice. This suggests that changes in serum bile acids are independent of de novo synthesis. We speculate that the increase of bile acids in the serum of GenX and PFOS exposed mice is due to changes in their disposition. It is known that PFAS, including PFOA and PFOS, inhibits the human sodium taurocholate cotransporting polypeptide (NTCP) (Ruggiero et al. 2021; Zhao et al. 2015). NTCP is one of the major bile acid influx transporters located on the basolateral sides of hepatocytes (Watashi et al. 2014). At 10 μM concentrations, PFOS act as inhibitors of NTCP, displacing bile acids, which in turn increases the serum levels of bile acids (Zhao et al. 2015). The serum concentrations for PFOS in our model was 8.96 μM, potentially in the NTCP inhibition range. Further studies are needed to determine the inhibitory effects of GenX on NTCP or other bile acid transporters, such as OATs and OATPs. Our data showed no significant increase in serum bile acids in PFOA treated mice. This could be partly due to the large variability in bile acid levels within the treatment group. As most PFAS studies have been on the effects of bile acid transporters *in vitro*, more *in vivo* experiments are needed to clarify the role of bile acid transporters in the deposition of PFAS.

### Induction of proliferation in response to PFAS exposure

PFAS induce cell proliferation in rodent livers through PPARα (Corton et al. 2018; Klaunig et al. 2003). However, because PPARα RNA and protein is expressed at higher levels in rodents compared to humans and has slightly different ligand binding domains of PPARα, relevance of PPARα activation as a mechanism has been questioned (Corton et al. 2014; Thomas et al. 2015; Wolf et al. 2008). Humanized mice treated with fenofibrate, a PPARα agonist, induced proliferation in only mouse hepatocytes and not human hepatocytes, indicating that proliferation induced by PFAS in humanized mice is PPARα-independent (Tateno et al. 2015) Previous studies from our group using primary human hepatocytes showed significant induction of the cell cycle gene *CCND1* by PFOA and PFOS (Beggs et al. 2016), and several promitogenic proteins, including Ki67 and CDK4 by GenX (Robarts et al. 2022b). The humanized mouse studies corroborated these findings and demonstrated that PFOA, PFOS, and GenX induced proliferation in human hepatocytes *in vivo*. We observed a significant increase in Ki67 and PCNA staining after PFOA and PFOS treatments, with a trend toward an increase following GenX exposure. This was accompanied by the induction of CCND1 protein expression in GenX, PFOA, and PFOS. Taken together, these data, along with previously published primary human hepatocyte exposures, show that PFOA, PFOS, and GenX induce PPARα independent hepatocyte proliferation at these occupationally relevant concentrations. Intriguingly, an increase in proliferation did not lead to an increase in the liver-weight-to-body-weight ratios of the humanized mice. One explanation could be a concomitant increase in cell death, both apoptosis and possibly necroptosis, after PFAS exposure. Our RNA-Seq data revealed significant activation of p53 in all PFAS exposed mice as compared to the control group, indicating a balance between cell proliferation and cell death. This suggests that PFAS could act as a promoter in hepatocellular carcinomas in hepatocytes possessing loss of function mutations in p53.

### Implications of changes in NR1D1 activity after PFAS exposure

One of the most intriguing findings of our studies is the induction of NR1D1, a nuclear receptor involved in the regulation of circadian rhythm, by all three PFAS. In general, light is the signal that starts clock synchronization across organs. The signal from the optic nerve travels to the suprachiasmatic nucleus (SCN) in the hypothalamus, which then communicates with the organs utilizing a variety of cellular and molecular cues, within the endocrine and nervous systems (Mukherji et al. 2019; Tahara and Shibata 2016). The orphan nuclear receptor NR1D1 (also known as Rev-Erbα), which functions as a transcriptional suppressor, regulates the intracellular circadian clock (Ramakrishnan and Muscat 2006). During the rest phase or night cycle, the two transcription factors BMAL1 and CLOCK heterodimerize, bind to the E-box promoter, and upregulate a plethora of genes, including proteins involved in the negative feedback loop, such as *PER1/2*, *CRY1/2*, *NR1D1*, and *NR1D2* (Mukherji et al. 2019). During the active phase or light cycle, NR1D1 binds to the E-box promoter of BMAL1, transcriptionally inhibiting BMAL1 production. This in turn decreases *NR1D1* mRNA levels until the night cycle when BMAL1 expression increases due to the loss of NR1D1(Mukherji et al. 2019). Dysregulation of this feedback loop is known to enhance liver disease progression, making it a critical KE in the AOP of hepatotoxicity (Mukherji et al. 2019; Tahara and Shibata 2016).

The effects of PFAS on circadian rhythm have never been documented. Our studies indicate that PFOA, PFOS, and GenX could disrupt circadian rhythm through the inhibition of NR1D1. We found an increase in E-box target genes, including a significant induction of *NR1D1* mRNA and protein. However, when measuring BMAL1 levels, there were very few differences between the treatments and control groups, indicating that NR1D1 activity is preferentially inhibited by these PFAS. Studies on determining cellular location, such as IHC and nuclear/cytoplasmic western blots, indicated that NR1D1 is heavily localized in the nucleus, indicating that PFAS do not interfere with nuclear translocation. However, *in silico* modeling suggests that PFOA, PFOS, and GenX all directly inhibit NR1D1 by binding to its DNA-binding domain. Because circadian rhythm regulates multiple processes, including xenobiotic absorption, distribution, metabolism, and excretion, and because its disruption exacerbates liver diseases, it is critical to understand whether PFAS are circadian rhythm disruptors (Baraldo 2008; Tahara and Shibata 2016). Future studies are necessary to investigate the full impact of PFAS on circadian rhythm utilizing human cells and humanized mice.

The humanized liver model comes with two main caveats. First, there may be a disruption in the cross talk between tissues/cells originating from mice and human hepatocytes. Apart from the hepatocytes, all other non-parenchymal cells and other organs are of mouse origin. The second limitation is that the hepatocytes used for repopulation are all derived from a single human donor, which can lead to individual bias.

In summary, our studies demonstrate that it is important to study PFAS in human-relevant models. These data indicate that humanized mice are an excellent model to identify human relevant AOPs of toxic exposure.

## Acknowledgements

The research support for this work was from NIH [grant numbers P20 RR021940-03 & S10 OD021743], NIH NIGMS [grant numbers P30 GM118247 & P30 GM122731], and NIH NIDDK [grant numbers NIH R01 DK0198414 & NIH R56 DK112768]. In addition, we thank Kaberi P Das for her technical support in the mass spectrometry analysis and Dr. Mark J. Strynar for measuring the PFAS in the serum and liver samples.

---

**Supplementary Figure 1.**
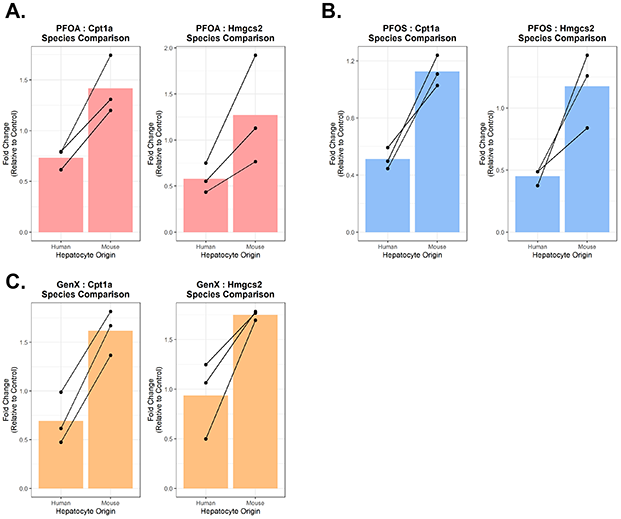
PFAS caused an induction of PPARα target genes expression in mouse hepatocytes and not in humans. qPCR of *Cpt1a/CPT1A* and *Hmgcs2/HMGCS2* using rodent and human specific primers in (A) PFOA, (B) PFOS, and (C) GenX treated humanized mice. Fold changes are relative to the untreated group. Gene expressions were normalized to the respective species housekeeping gene (*Gapdh/GAPDH*). Connecting lines represent expression of mouse and human genes derived from the same liver with bars representing the average fold change.

**Supplementary Figure 2.**
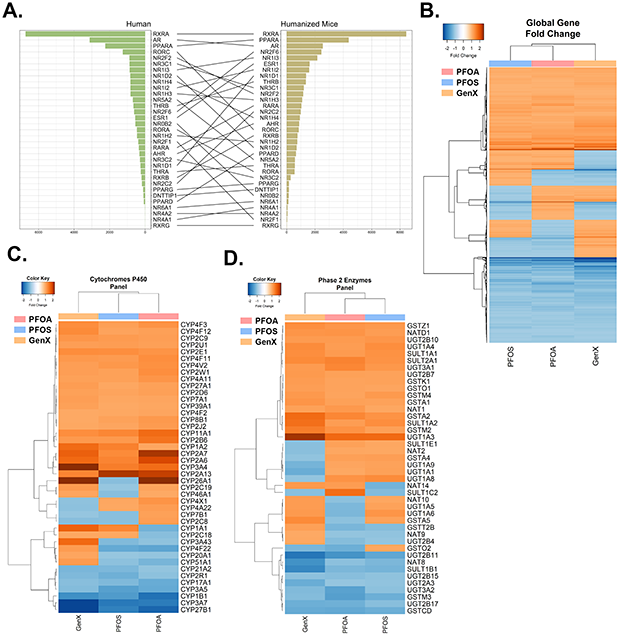
Global changes of RNA-Seq of humanized mice treated with PFOA, PFOS, GenX. (A) Connecting bar plot of the expression level of ligand-binding nuclear receptors in livers of human and control humanized mice, where the y-axis is the gene, and the x-axis is the normalized DESeq2 normalized counts. Expression levels were ranked from highest to lowest, with the black line linking the gene of each species. Heatmap of (B) global gene expression, (C) cytochrome P450 metabolizing enzymes, and (D) phase 2 enzymes. Blue and orange represent negative and positive fold changes, respectively.

**Supplementary Figure 3.**
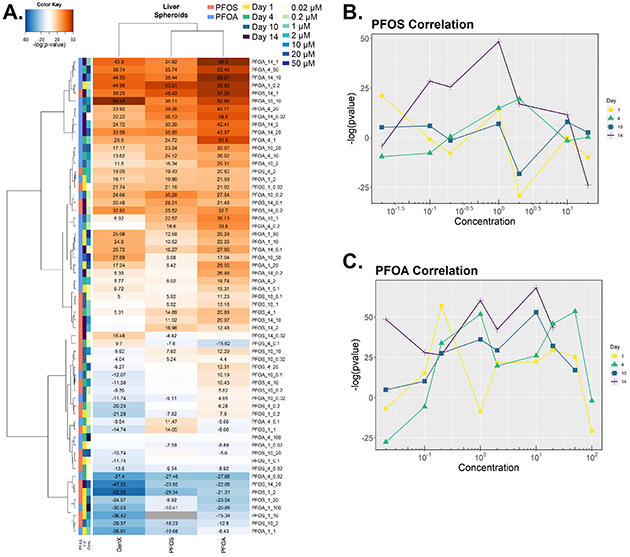
RNA-Sequencing of PFOA and PFOS treated humanized mice is significantly correlated with human hepatic spheroids exposed to PFOS and PFOA for 14 days. (A) Heatmap of -log(p-values) from BSCE comparing the DEGs from exposed human hepatic spheroids and the treated humanized mice. Blue and orange represent negative and positive correlations, respectively, and no assigned p-value is represented by gray. Column colors represent the compound (PFOA or PFOS) the spheroids were exposed to, the exposure time (1, 4, 10, 14 days), and the concentration of the compound (0.02, 0.2, 1, 2, 10, 20, 50 μM). Each number in the cells of the heatmap represents the significant -log(p-value)s with no number indicating no significance. Line graphs of each -log(p-value) for (B) PFOS and (C) PFOA exposed human hepatic spheroids. Where color and shape indicate the length of spheroid exposure, the y-axis represents the - log(p-value) and the x-axis represents the concentration (μM) exposed to the spheroids.

**Supplementary Figure 4.**
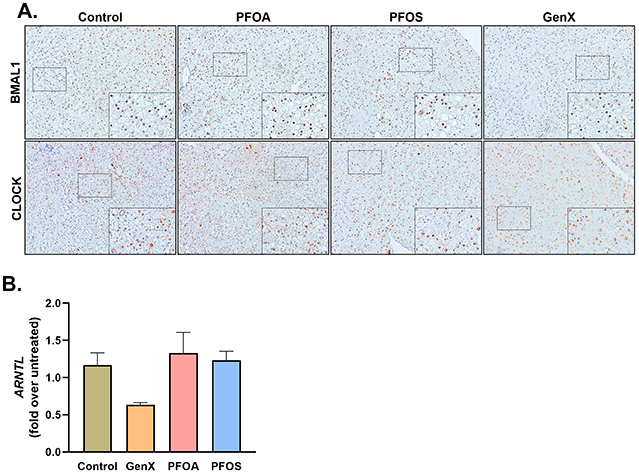
PFAS caused no change in BMAL1 and CLOCK nuclear localization. Photomicrographs of liver IHC of BMAL1 and CLOCK for control and PFAS treated humanized mice at 200x. The small box corresponds to the 400x magnification image. qPCR of *ARNTL* (BMAL1 gene), where samples were first normalized to the 18s housekeeping gene and then to the control group. Bars represent mean ± SEM.

